# Custom hereditary breast cancer gene panel selectively amplifies target genes for reliable variant calling

**DOI:** 10.1101/322180

**Authors:** Setor Amuzu, Timothée Revil, William D. Foulkes, Jiannis Ragoussis

## Abstract

**Background:** Target enrichment coupled with next generation sequencing provide high-throughput approaches for screening several genes of interest. These approaches facilitate screening a panel of genes for mutations associated with inherited breast cancer for research, diagnostic, and genetic counseling applications.

**Objective:** To evaluate the performance of our custom 13 gene breast cancer panel, based on singleplex PCR, developed by WaferGen BioSystems. The panel was evaluated using patient-derived DNA samples, in terms of target enrichment efficiency, off-target enrichment, uniformity of target capture, effect of GC content of target regions on coverage depth, and concordance with validated variant calls.

**Results:** At least 90% of target sequence for each gene was captured at 30x or greater. We evaluated uniformity of target capture across samples by calculating the percentage of samples with at least 90% of total target captured at 100x or greater and found 92% (33/36 samples) uniformity for our panel. Off-target enrichment ranges between 7.2% and 22.3%. We found perfect concordance between our custom panel and the Qiagen human breast cancer panel for functionally annotated variant calls in high read depth shared target regions. Altogether, there was agreement between the panels for 779 variants at 41 loci. We also confirmed 10 pathogenic mutations, initially discovered by Sanger sequencing, in the appropriate samples following target enrichment using our custom WaferGen panel.

**Conclusion:** Our custom hereditary breast cancer panel is sensitive to the desired target genes and facilitates deep sequencing for reliable variant calling.

## Introduction

Targeted sequencing of genomic regions of interest remains a viable way to study diseases with known genetic associations and could be routinely used in molecular diagnoses. It is cheaper, produces data that is more tractable, substantially faster to analyze, and more easily understood for functional interpretation compared to whole exome sequencing. Investigating a panel of multiple genes can inform clinicians and researchers of genetic variants that may be associated with breast cancer risk (Easton et al., 2015; Kurian et al., 2014; Walsh et al., 2010). Breast cancer is the most prevalent cancer among women, and one of the leading causes of cancer deaths among women. In 2013, 1.8 million women died from breast cancer worldwide (Global Burden of Disease Cancer Collaboration et al., 2015). A number of genes of varying penetrance have been associated with susceptibility to breast cancer (Michailidou et al., 2013; Rahman et al., 2007; Wooster et al., 1995). Breast cancer gene panel testing is now commercially available for molecular biology applications, with a variety of pre-designed and customizable panels on offer. In this study we evaluated a custom panel designed for the SmartChip approach by WaferGen Biosystems Inc. (now acquired by Takara Bio USA Holdings, Inc.). Although the SmartChip has been evaluated using DNA derived from cancer cell lines (De Wilde et al., 2014), to our knowledge an evaluation for its application in inherited breast cancer still needs to be presented. We designed our panel to capture the entire coding sequence of 13 genes frequently mutated in inherited breast cancer cases, representing a target of 150Kb in total. Our panel includes nine established breast cancer susceptibility genes with high and moderate risk. For these genes *(BRCA1, BRCA2, ATM, CHEK2, TP53, CDH1, STK11, PALB2, PTEN)* association with breast cancer risk has been established, primarily, on protein truncating variants (Easton et al., 2015). Low risk or proposed breast cancer susceptibility genes *BRIP1, RAD51C, RAD51D,* and *PPM1D* were also included in spite of their relatively low mutation rate in inherited breast cancer cases (Slavin et al., 2017). All but two *(RAD51C, RAD51D)* of the genes on our panel are captured in version 84 of the COSMIC cancer gene census list (Futreal et al., 2004). A summarized description of target genes for our panel is presented in Table 1.

**Table 1.** List of 13 target genes for our hereditary breast cancer panel and their associated cancer risks. TSG – tumor suppressor gene. *Risks mainly associated with biallelic pathogenic variants.

| Gene symbol | Name | Major associated cancer risk | Summary of function | Role in cancer | References (PubMed IDs) |
| --- | --- | --- | --- | --- | --- |
| *ATM* | Ataxia telangiectasia mutated | Breast, leukemia, lymphoma | DNA double- strand break repair | TSG | 15928302, 21787400, 16832357 |
| *BRCA1* | Familial breast/ ovarian cancer gene 1 | Breast, ovarian | Homologous recombination DNA repair, replication fork repair | TSG | 2270482, 7545954, 7939630 |
| *BRCA2* | Familial breast/ovarian cancer gene 2 | Breast, ovarian, pancreatic | Homologous recombination DNA repair | TSG | 8524414, 10433620, 8640235 |
| *BRIP1* | BRCA1 interacting protein C-terminal helicase 1 | Breast | Homologous recombination DNA repair | TSG | 16959974, 17033622 |
| *CDH1* | Cadherin 1, type 1, E-cadherin (epithelial) | Breast, gastric | Calcium- dependent cell adhesion receptor | TSG | 17660459, 7961105 |
| *CHEK2* | Checkpoint Kinase 2 | Breast, thyroid, prostate | DNA double-strand break repair | TSG | 27751358, 16880452, 25583358, 12533788 |
| *PALB2* | Partner and localizer of BRCA2 | Breast, Wilms tumor*, medulloblastoma* | Homologous recombination DNA repair | TSG | 17200668, 19812674 |
| *PPM1D* | Protein phosphatase, Mg<sup>2+</sup>/Mn<sup>2+</sup> dependent 1D | Breast, ovarian, glioma | Negative regulator of *TP53* and *CHEK2* | Oncogene | 12021784, 23242139, 23649806 |
| *PTEN* | Phosphatase and tensin homolog gene | Breast, prostate, endometrial, lung | Modulator of PI3K/AKT pathway, regulates p53 | TSG, oncogene | 9439675, 9140396, 12620407 |
| *RAD51C* | RAD51 paralog C | Ovarian | Homologous recombination DNA repair, genome stability | TSG | 20400964 |
| *RAD51D* | RAD51 paralog D | Ovarian | Homologous recombination DNA repair, genome stability | TSG | 22415235, 22652533, 28656019 |
| *STK11* | Serine/threonine kinase 11 gene | Breast, ovarian, testicular, pancreatic | Regulates cell polarity | TSG | 16707622, 10429654 |
| *TP53* | Tumor protein p53 | Breast, glioma, leukemia, lung, and other pediatric- and adult-onset cancers | Coordinates cellular response to DNA damage, hypoxia, and other stress signals | TSG, oncogene | 2052583, 11219776, 11879567, 10427138, 1656362 |

PCR-based target enrichment (TE) is commonly used to discover novel variants in known breast cancer susceptibility genes or validate mutations such as single nucleotide polymorphisms (SNPs) and small insertions and deletions (indels). Singleplex PCR strategies have been found to detect mutations present at less than 5% frequency in tumor samples (Kozarewa et al., 2015). Our custom panel is based on WaferGen’s Seq-Ready™ TE Custom DNA Panel. This panel relies on WaferGen’s high-density SmartChip™ TE singleplex PCR technology to capture up to 2.5 Mb of cumulative sequence using massively parallel singleplex PCR (WaferGen Press, 2015). Each target region is amplified in an individual PCR reaction in a single well on the chip. This design is intended to improve sensitivity and specificity, and reduce primer-primer interaction as well as other interference often associated with multiplex PCR. WaferGen expects this approach to enable more reliable variant calling and obviate the need for variant validation by Sanger sequencing, for example. They report 84% mutation validation rate, representing 21 out of 25 mutations, based on sequencing data from 8 genes in a study using 15 cancer cell lines (De Wilde et al., 2014). Mutations validated in this study include SNPs and indels. Additionally, primers are designed such that no known SNPs with population frequency greater than 0.5% are included in last 10 nucleotides on the 3’ end of a primer, to minimize off-target hybridization and reduce risk of allelic dropouts (De Wilde et al., 2014). Amplicons are designed to span beyond target exons and overlap each other to increase the probability of capturing target regions (Figure 1).

**Figure 1.**
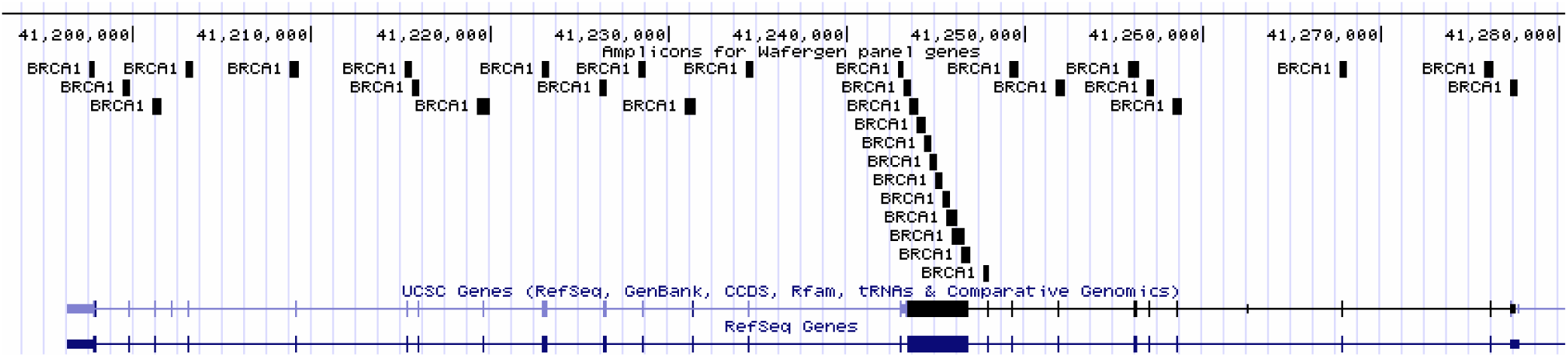
*BRCA1* amplicons of Wafergen panel are mapped to *BRCA1* exons of GRCh37/hg19 using UCSC genome browser (Kent et al., 2002). Amplicons overlap target exons to maximize capture. Amplicons are designed to span exon-intron boundaries and untranslated regions.

To develop our custom breast cancer panel, we provided WaferGen with coordinates of our genes of interest, desired amplicon length, sequencing platform, and samples. Genomic coordinates for our WaferGen panel is provided as Supplementary file 1. Our custom breast cancer panel targets 314 regions representing all exons for 13 genes which are listed, along with other panel features, in Table 2.

**Table 2.** Target genes of custom WaferGen hereditary breast cancer panel. Panel is based on Seq-Ready™ TE MultiSample Custom DNA Panel (Part number: 440-000053) product line. Amplicon size ranges from 350 – 749 bases. Capture and exon coordinates are based on human reference genome assembly GRCh37/hg19.

| Gene (RefSeq accession) | Number of exons | Number of amplicons | Exon target size (bp) | Capture size (bp) |
| --- | --- | --- | --- | --- |
| *ATM*<br>(NM_000051) | 63 | 72 | 13,210 | 32,303 |
| *BRCA1*<br>(NM_007300) | 24 | 33 | 6,083 | 15,820 |
| *BRCA2*<br>(NM_000059) | 27 | 42 | 11,002 | 21,744 |
| *BRIP1*<br>(NM_032043) | 20 | 28 | 6,512 | 13,454 |
| *CDH1*<br>(NM_004360) | 16 | 21 | 4,831 | 9,712 |
| *CHEK2*<br>(NM_001005735) | 16 | 16 | 2,003 | 8,109 |
| *PALB2*<br>(NM_024675) | 13 | 18 | 4,071 | 8,255 |
| *PPM1D*<br>(NM_003620) | 6 | 10 | 3,040 | 4,553 |
| *PTEN*<br>(NM_000314) | 9 | 30 | 5,556 | 14,728 |
| *RAD51C*<br>(NM_058216) | 9 | 9 | 1,291 | 3,844 |
| *RAD51D*<br>(NM_002878) | 10 | 9 | 1,461 | 4,448 |
| *STK11*<br>(NM_000455) | 10 | 15 | 3,286 | 7,596 |
| *TP53*<br>(NM_001276760) | 11 | 11 | 2,602 | 6,079 |
| **Total** | 234 | 314 | 64,948 | 150,645 |

In this study, our goals were to assess how well our custom panel captures its target genes, examine the effect of GC content of target regions on coverage depth, and measure concordance in genotype calls between our panel and known variant calls obtained from Sanger sequencing and target enrichment using the Qiagen human breast cancer panel.

## Materials and Methods

### Samples

Peripheral blood samples from 36 hereditary breast cancer cases were selected for testing on our custom WaferGen breast cancer panel. A subset (19) of these samples were tested using the validation panel, Qiagen Human breast cancer panel. The Human Breast Cancer GeneRead DNAseq Gene Panel targets hotspots in 20 genes commonly mutated in human breast cancer samples using a multiplexed PCR assay. The Qiagen panel is compatible with both germline and tumor mutation profiling and shares a common target region with our custom panel. Genomic DNA (gDNA) was extracted from blood samples and quantified using Qubit dsDNA HS.

### Target enrichment and library preparation

Purified genomic DNA from our Qiagen samples were subjected to target enrichment using the GeneRead DNAseq Targeted V2 Panel and the GeneRead DNAseq PCR Kit V2 (comprising oligonucleotides, enzymes and buffers) according to manufacturer’s instructions. Wafergen’s SmartChip target enrichment platform, previously described by De Wilde *et al.* (De Wilde *et al.,* 2013), was used to amplify our custom genomic targets. Reaction mix for WaferGen samples was prepared using 350ng of gDNA and KAPA2G DNA polymerase. Reaction volume per well of SmartChip was 50nl. PCR was performed using a WaferGen-modified thermal cycler and resulting amplicons pooled by centrifugation.

Nextera XT tagmentation protocol was used to prepare the sequencing library from WaferGen amplicons while the ligation-based NEBNext method for Illumina was used to prepare the sequencing library from Qiagen amplicons. WaferGen amplicons, due to their size (350 – 749 bases), were fragmented prior to ligating read-specific sequencing primers, index, and adapter sequences. However, there was no fragmentation step in the library preparation for Qiagen samples because these amplicons are smaller (48 – 143 bases).

### Sequencing

Samples were sequenced by the McGill University and Génome Québec Innovation Centre sequencing service. All samples were sequenced on the Illumina HiSeq 2500 to produce 150bp, paired-end reads. Samples were sequenced in rapid run mode.

### Data analysis

Raw sequencing data was filtered using Trimmomatic (Bolger et al., 2014) with Phred Quality score of 20 as cutoff. Filtered sequence reads for each sample were aligned to human reference genome build GRCh37/hg19 using BWA-MEM (Li and Durbin, 2010). The alignment files were converted from SAM to BAM format and sorted. BEDtools (Quinlan and Hall, 2010), and bamstats05 from the Jvarkit (Pierre, 2015) package were used to determine breadth and depth of coverage metrics for individual target intervals using each sample’s BAM file. Coverage metrics for samples were aggregated for each panel using custom R scripts. Plots and statistics were also generated using R programming language for statistical computing (R Core Team, 2016). Variant calling was done using HaplotypeCaller of GATK version 3.7-0-gcfedb67 (McKenna et al., 2010), according to best practices. Variants for both sets of samples were called as a cohort. The resulting VCF file was annotated with dbSNP (Sherry et al., 2001) identifiers from dbSNP146 using SnpSift (Cingolani et al., 2012). Variants were normalized using bcftools, and partial genotypes were converted to complete genotypes using genotype likelihood scores. Annotated VCF was filtered with the vcfR (Knaus and Grunwald, 2016) package and custom R scripts. Variants were filtered using total coverage threshold of 30. Only variants with dbSNP identifiers (rs IDs) were considered. For the Qiagen panel, only variants outside of the target enrichment primer regions were considered for assessment of concordance with our custom panel since variants near the amplicon boundaries can cause misalignments of multiple reads leading to false-positive or false-negative variant calls (Vijaya Satya et al., 2014). SnpSift Concordance was used to measure genotype concordance. Integrative Genomics Viewer (Thorvaldsdóttir et al., 2013) was used to manually inspect some variant calls.

## Results and Discussion

### Target Enrichment Efficiency

Target capture was inferred from sequence reads, derived from target enrichment amplicons, that mapped to target genes. We defined coverage breadth as the proportion of unique reads that were unambiguously mapped to target regions, and determined coverage breadth per gene and per sample at coverage depth thresholds of 1x, 30x, 50x, and 100x. By examining breadth of coverage per gene and per sample we assessed the sensitivity of capture for each target gene, and uniformity of total target enrichment across samples. In Figure 2, we show that more than 90% of target sequence for each gene is captured at 30x or greater. The majority of exons are captured at reasonable depth for germline variant calling.

Similarly, coverage depth for target exons across samples is also adequate for germline variant calling with 97% (35/36) of samples captured at least 30x (Figure 3). Uniformity of target capture across samples is high – 92% of samples (33/36) capture 90% of target with coverage depth of at least 100x.

**Figure 2.**
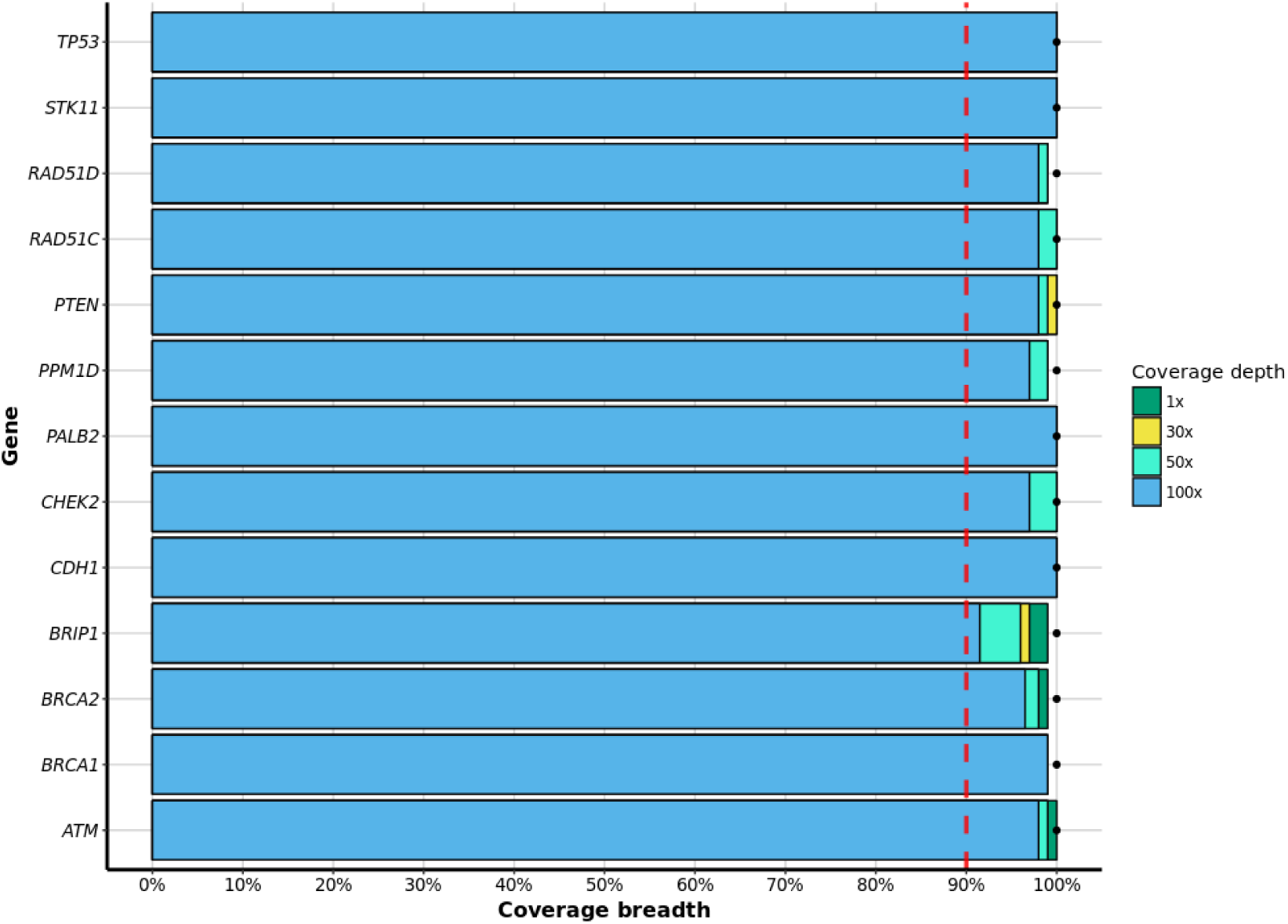
A barplot showing the median percentage of target genes captured at 1x, 30x, 50x and 100x coverage depth thresholds. The red dashed line at 90% on the Coverage breadth axis represents our expected minimum breadth of coverage for each gene. The black dot beside each bar denotes the maximum breadth of coverage at 1x for a specific target gene.

**Figure 3.**
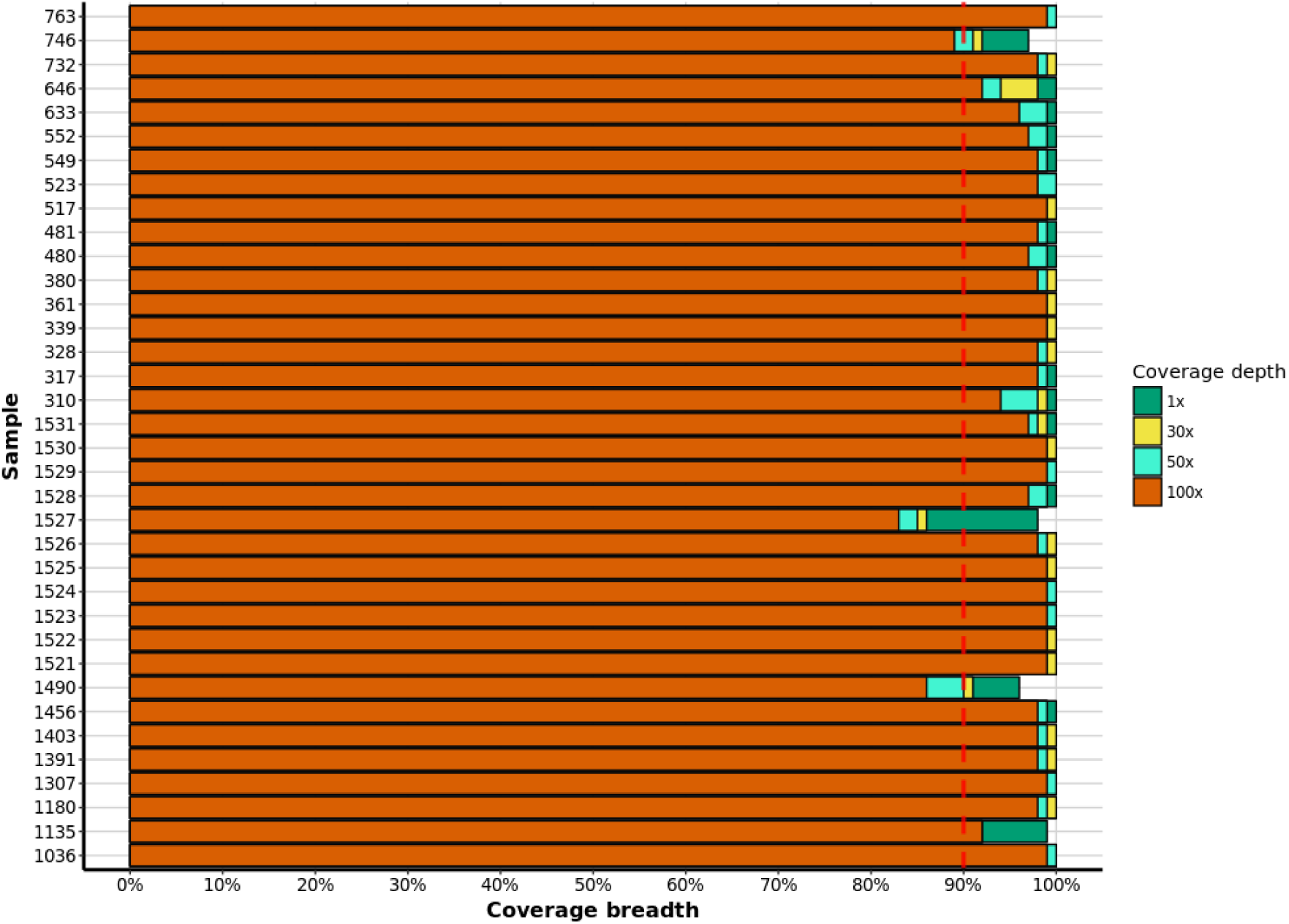
Median percentage of target exons captured for each sample at 1x, 30x, 50x and 100x coverage depth thresholds. The red dashed line at 90% on the Coverage breadth axis represents our minimum expected breadth of coverage for each sample.

We assessed the effect of GC content on coverage depth of amplicons. The Wafergen panel is capable of enriching high GC (> 70%) and low GC (< 30%) regions, but peak coverage depth lies between these extremes of GC as expected from Illumina sequencing (Figure 4). Coverage depth tends to increase with GC content. This is driven by the sharp increase in coverage depth from low GC (30% - 40%) amplicons to moderate GC (45% - 65%) amplicons, and does not represent overall increase in coverage depth across the entire spectrum (26% - 81%) of GC content of amplicons.

**Figure 4.**
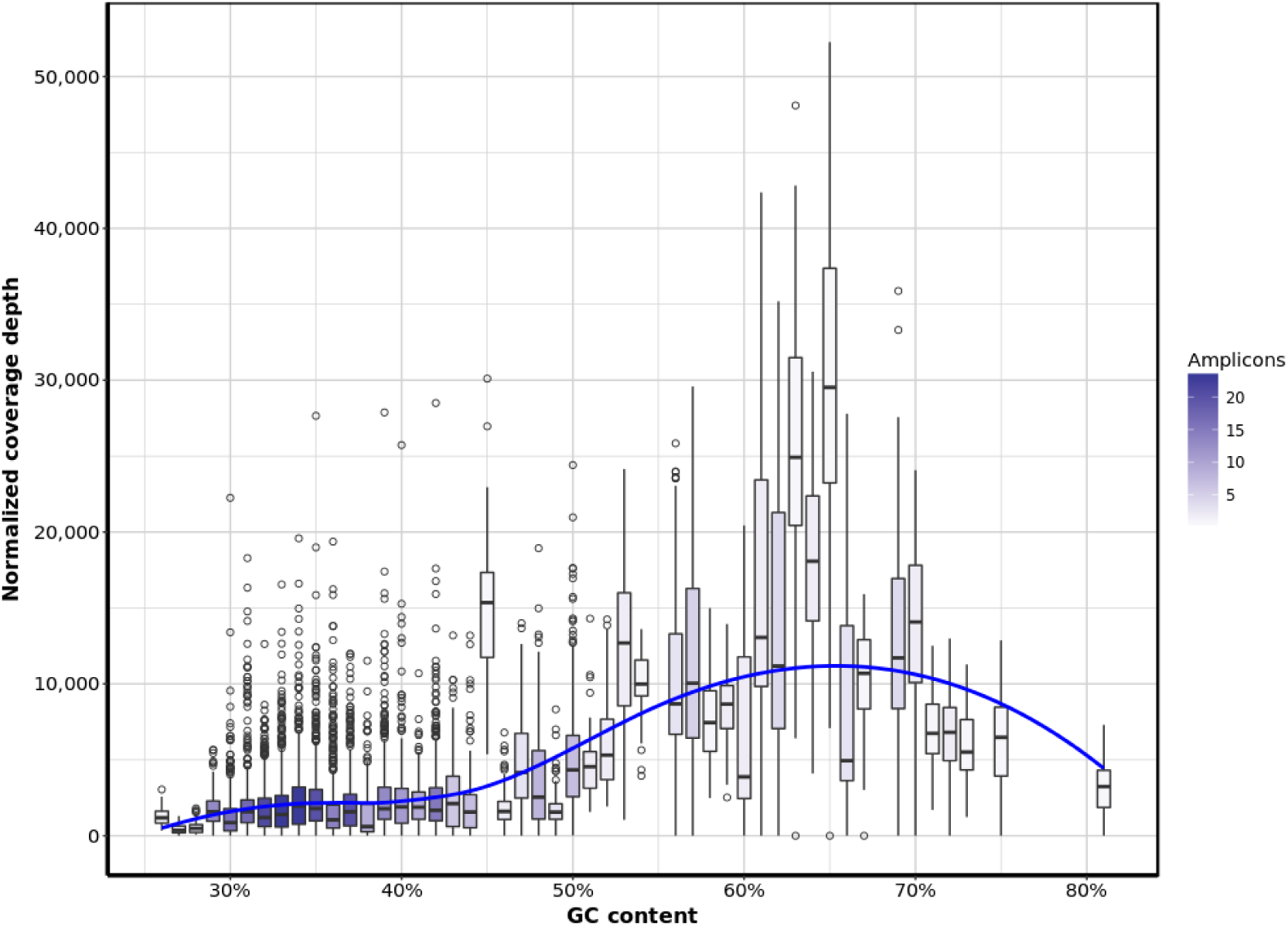
Relationship between GC content of amplicons and coverage depth. Normalized coverage depth is on-target reads per kilobase target region, million mapped reads and number of overlapping amplicons. Amplification of target occurs across the width of GC content (26% - 81%).

### Off-target enrichment

We define off-target enrichment as the percentage of reads mapping to other regions of the genome outside the capture target per sample. We observed that off-target enrichment ranges between 7.2% and 22.3%, with mean of 17.3% and standard deviation 3.7%.

### Concordance in genotype calls

Using sanger sequencing, 10 germline heterozygous, pathogenic mutations were identified for 10 WaferGen samples. These mutations were confirmed in the relevant samples with the same zygosity identified by Sanger sequencing (Table 3).

**Table 3.** List of mutations validated by Sanger sequencing for 10 WaferGen samples. Mutations include SNPs and short deletions.

| Gene | Variant | dbSNP ID | Sample |
| --- | --- | --- | --- |
| *BRCA1* | c.68_69delAG (p.Glu23fs) | rs386833395 | 1521 |
| *BRCA2* | c.5857G>T (p.Glu1953*) | rs80358814 | 1522 |
| *TP53* | c.730G>A (p.Gly244Ser) |  | 1523 |
| *ATM* | c.2572T>C (p.Phe858Leu) | rs1800056 | 1524 |
| *PALB2* | c.229delT (p.Cys77fs) | rs180177084 | 1525 |
| *CHEK2* | c.1100delC (p.Thr367fs) | rs555607708 | 1526 |
| *STK11* | c.876delC (p.Tyr292fs) |  | 1527 |
| *PTEN* | c.389G>A (p.Arg130Gln) | rs121909229 | 1528 |
| *CDH1* | c.382delC (p.His128fs) |  | 1529 |
| *PPM1D* | c.1654C>T (p.Arg552*) |  | 1530 |

We also assessed concordance in genotype calls between our custom WaferGen panel and Qiagen’s human breast cancer panel over a shared target region of 33Kb, comprising amplicons for *TP53, ATM, BRCA1, BRCA2, PTEN,* and *CDH1.* A subset of 19 samples were available for this assessment. We considered genotypes that were called with read depth of 30 or greater and variants that were recorded in dbSNP. In Table 4, we present the frequencies of genotypes that meet these conditions. We found perfect concordance between genotype calls for the Wafergen and Qiagen panels over a shared target region for the same samples at high confidence, annotated variant loci.

**Table 4.**
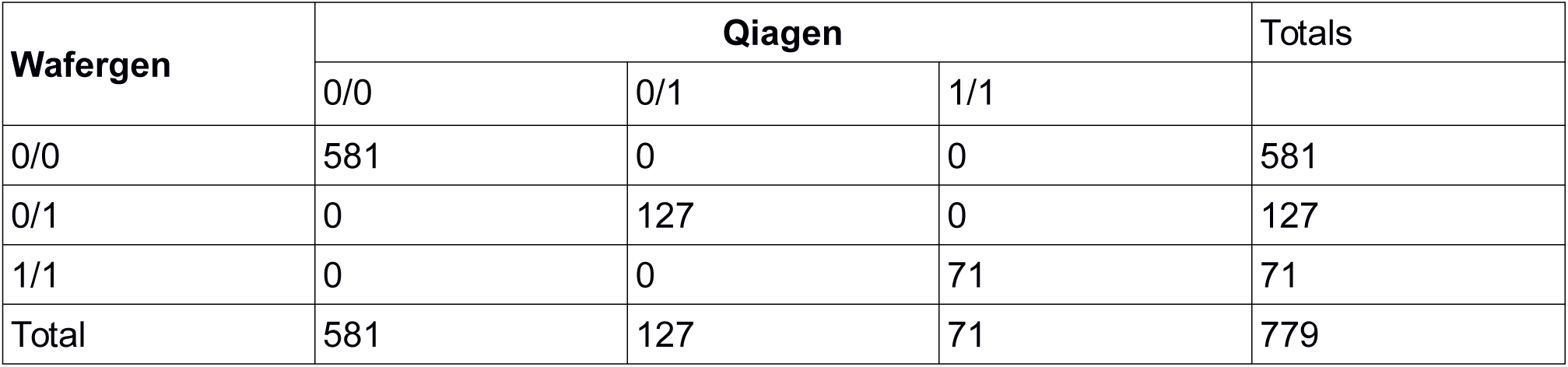
Contigency table of genotype counts for 41 variant loci across 19 matching samples representing the relationship between genotype calls for WaferGen and Qiagen panels. 0/0 is homozygous reference, 0/1 is heterozygous, and 1/1 denotes homozygous alternate.

## Conclusions

We demonstrated that our custom hereditary breast cancer panel adequately captures all exons of 13 breast cancer susceptibility genes for next generation sequencing and subsequent detection of germline mutations associated with breast cancer. The WaferGen panel is sensitive for target genes with minimal off-target amplification. At least 90% of target region is captured at reasonable depth (at least 30x) for reliable variant calling, with uniform capture across samples. Additionally, our custom panel can confirm known pathogenic mutations, including SNPs and short deletions, and is comparable to more established Qiagen human breast cancer panel in terms of amplifying exome targets for high confidence variant calling.

## Author contributions

WDF and JR conceived the study and critically revised the mauscript. WDF arranged the procurement of samples. JR coordinated sequencing and target enrichment. JR and WDF were involved in selecting genomic coordinates for the panel.

TR performed target enrichment, and data analysis (quality control, sequence alignment, and variant calling). TR also revised the manuscript.

SA performed data analysis (coverage analysis, variant concordance assessment) and visualization, and prepared the first draft of this manuscript.

## Acknowledgements

We are grateful to Nelly Sabbaghian and Evan Weber for preparing samples.

